# Bigger Isn’t Always Better: Comparing System Size, Hydration, and Software for Lipid Membrane Analysis

**DOI:** 10.64898/2026.01.20.700632

**Authors:** F. Carvalho, P. Maximiano, P.N. Simões, M. Hashemi

## Abstract

The structural and dynamic properties of membranes are known to vary with bilayer size and hydration. While Molecular Dynamic (MD) simulations are a powerful tool for studying cellular membrane systems, the results can be sensitive to the analysis workflow and software. In this study, all-atom MD simulations (500 ns) were conducted on systems of 256, 512, and 1024 POPC lipids at 40, 80, and 160 waters per lipid. With these simulations, a two-fold study was performed: (1) to assess the convergence of structural and dynamic properties of POPC bilayers as a function of membrane size and hydration level using CPPTRAJ (CPP), including area per lipid (APL), bilayer thickness, order parameter, headgroup orientation, and lateral diffusion, and (2) to compare the analysis output and performance of four software packages: CPP, GROMACS (GRO), MDAnalysis (MDA), and LiPyphilic (LiP). For the first objective, our results show that the average values of the bilayer thickness, order parameter, and headgroup orientation are largely independent of the size and hydration levels studied. In contrast, lateral diffusion coefficient was sensitive to both size and hydration. We found that increasing the system size primarily decreased the statistical variance of the APL and thickness. For the second objective, all four packages produced consistent results for APL and thickness, with the most significant discrepancy being a known artifact from the `gmx order` tool when applied to unsaturated carbons. Performance bench-marks identified `CPP` as the fastest serial tool for all properties, whereas parallelization benefited `MDA` and `LiP` in some metrics. These findings provide a practical roadmap, demonstrating that moderately sized systems (e.g., 256L), combined with an optimized tool such as `CPP`, offer an efficient workflow for membrane structural property analysis.

## Introduction

Molecular dynamics (MD) simulations have become a powerful tool for investigating cellular membranes, which are dynamic and complex assemblies of lipids, proteins, and other molecules^1,2^. MD simulations are applicable to a wide range of membrane systems, from simple lipid mixtures to complex biological membranes that include, for example, proteins, glycolipids, sphingolipids, and ceramides^3^. However, accurately characterizing membrane properties from these simulations is non-trivial. The results are sensitive to physical parameters, such as membrane size and hydration level, and can also be influenced by the choice of analysis software.

Castro-Román et al. ^4^ investigated discrepancies between experimental and computational data for membrane properties. By comparing systems with 72 and 288 lipids at low hydration (5.4 waters/lipid), they found that the structural properties, such as the area per lipid (APL), order parameters, and density profiles, were largely independent of the system size. They concluded that the observed discrepancies were not due to artifacts of finite-size effects but rather due to other factors, such as the selected force field. The authors also reported that correctly modeling hydration is critical, as the APL is influenced by hydration, particularly in 1,2-dioleoyl-sn-glycero-3-phosphocholine (DOPC) bilayers. Similarly, Klauda et al.^5^ demonstrated that a system of 72 1,2-dipalmitoyl-sn-glycero-3-phosphocholine (DPPC) lipids was sufficient to calculate structural properties such as density profiles and order parameters. Other metrics, including the P– N dipole orientation, radial distribution functions, lipid rotational correlation functions, and C – H bond isomerization dynamics, were also independent of the system size, whereas lateral diffusion exhibited a clear size dependence on the system size.

Smith et al.^6^ emphasized that APL measurements in large membrane systems must account for membrane undulations or curvature, which are size-dependent phenomena. These undulations influence both the APL and membrane thickness, which are interrelated structural properties of the membranes. Farhangian et al. ^7^ observed that the APL increased with the hydration number, reaching a plateau at approximately 32 water molecules per lipid molecule. This increase was more pronounced in 1,2-dioleoyl-sn-glycero-3-phospho-ethanolamine (DOPE) and 1-palmitoyl-2-oleoyl-sn-glycero-3-phosphoethanolamine (POPE) than in DOPC and 1-palmitoyl-2-oleoyl-sn-glycero-3-phosphocholine (POPC), suggesting a correlation with the greater hydrophilicity of the phosphoethanolamine (PE) headgroup.

Högberg and Lyubartsev^8^ investigated the effects of hydration and temperature on the structural and dynamic properties of dimyristoylphosphatidylcholine (DMPC) bilayers using hydration levels of 5, 10, and 29 water molecules per lipid. At 30 °C, they observed that APL increased with hydration, while both lipid headgroup orientation angle and membrane thickness decreased. The dynamic properties, including the diffusion coefficients of both DMPC and water, exhibited a direct correlation with the hydration level, whereas the relaxation times of the lipids, head groups, and water decreased with increasing hydration.

While these studies have independently investigated system size^4–6^ and hydration levels^4,7,8^, their combined and interdependent effects on parameter convergence, particularly at the large system sizes and high hydration levels now common in modern simulations, are not fully characterized. Understanding this convergence is key to the efficient use of computational resources when selecting the system size and hydration level for simulations. Another important aspect of MD simulations is the analysis tools used to quantify the results obtained. In terms of analysis tools, CPPTRAJ (`CPP`) is the primary analysis package of the AmberTools suite, written in C++ for high performance and designed to run through its own scripting language^9^. Its core strength lies in its processing speed, making it an ideal choice for handling large trajectories on high-performance computing clusters. A wide range of standard biomolecular analyses, together with an extensive library of pre-written commands, contribute to its broad applicability^10^. The trade-off is reduced flexibility, a fixed command set, and syntax inconsistencies between versions. These factors can lead to a steep learning curve, particularly for complex analyses.

GROMACS (`GRO`) includes a powerful set of command-line tools bundled directly with the simulation engine. Its primary advantage is its full compatibility with the native GROMACS file formats (`.tpr`, `.xtc`, `.edr`), avoiding the parsing issues that often challenge external tools^11,12^. Its main limitation lies in performing complex analyses that require chaining multiple commands, which can only be achieved through shell scripting.

MDAnalysis (`MDA`) is an open-source Python library that adopts an object-oriented approach to simulation analysis^13^. Its main advantage is flexibility: by representing simulations as Python objects and leveraging the broader Python ecosystem —especially the NumPy and SciPy libraries —it enables virtually unlimited custom analyses. Its `MDA.Universe` parser can read data from almost any simulation package, making it a versatile tool for cross-software and comparative analysis^14^. To fully exploit the capabilities of this software, Python programming skills are required to build analyses from its fundamental components. The core functionalities of `MDA` and NumPy are described in detail by Michaud-Agrawal et al.^13^ and Gowers et al.^14^

LiPyphilic (`LiP`) is a Python library built directly on top of MDAnalysis and is specifically developed for lipid membrane analysis^15^. It offers high-level commands while retaining the flexibility of the Python environment, simplifying complex membrane-specific tasks—such as APL or order parameter analysis—into single, user-friendly functions. Its main limitation is its specialized scope; it is not intended for general molecular analysis. Although its feature set is growing, it remains less comprehensive than broader packages, such as CPPTRAJ or MDAnalysis.

The methodological question of how the choice of analysis software affects the characterization of simulation physical parameters, such as system size and hydration, has largely remained unaddressed. This question has practical implications because the selected software package can directly impact accuracy, computational demands, and analysis workflow.

Therefore, this study had two objectives. First, we aimed to assess the convergence of the structural and dynamic properties of POPC bilayers as a function of the membrane size and hydration level. Second, we compared the numerical results and performance of different software packages for analyzing these properties.

To address these objectives, we performed all-atom molecular dynamics simulations on lipid bilayer systems composed of 256, 512, and 1024 POPC lipid molecules, each size simulated at three different hydration levels (40, 80, and 160 water molecules per lipid), with each system simulated in triplicate. Key equilibrium structural metrics, namely the APL, bilayer thickness, lipid order parameter, and lipid headgroup orientation, as well as a dynamic metric, namely the lateral diffusion coefficient, were evaluated. For the first objective, `CPP` was selected due to its high performance and extensive command libraries. For the second objective, each property was computed using `CPP`, `GRO`, `MDA`, and `LiP`, and the results were systematically compared and discussed to assess the consistency, reliability, and methodological differences across these platforms.

## Methods

Three independent all-atom molecular dynamics (MD) simulations of 500 ns starting from the same initial coordinates were performed for each *i*L–*j*W membrane system, where *i* = 256, 512, 1024 lipid molecules and *j* = 40, 80, 160 water molecules per lipid. The simulation boxes were constructed with the aid of the CHARMM-GUI^16^. The initial box sizes and compositions of the simulated systems are listed in Table SI.1 (Supporting Information). All simulations were performed under periodic boundary conditions in the NPT ensemble at 300 K, with temperature controlled via a Langevin thermostat (collision frequency: 1.0 ps^−1^)^17^, and pressure maintained at 1.0 bar using a semi-isotropic Monte Carlo barostat with a pressure relaxation time of 1.0 ps^18^. An integration time step of 2 fs was used throughout the study. Long-range electrostatic interactions were treated using the Particle Mesh Ewald (PME) method, whereas a short-range cutoff distance of 12.0 Å was employed for all nonbonded interactions^19^. The bond lengths involving hydrogen atoms were constrained using the SHAKE algorithm^20^.

A crucial factor when performing biologically relevant MD simulations is the choice of force field. To faithfully capture the dynamics of a system, the force fields for each biomolecular component (e.g., lipids, proteins, and nucleic acids) must be mutually compatible, ensuring that all interactions among different molecule types are accurately described and validated. In this study, we used the Amber Lipid 17 force field^21^ and the TIP3P water model^22^. This choice was motivated by our use of the Amber family of protein force fields^23^, as well as our intended use of Amber nucleic-acid force fields^24^ in combination with lipids. Stockholm Lipids^25^ is another lipid force field compatible with the Amber family; however, its lipid coverage is more limited compared to Amber Lipids 17. CHARMM is a separate family of force fields with a very extensive parameter set, covering numerous biomolecules and their combinations.

### Software and Hardware Specifications

CPPTRAJ v6.18.1 (`CPP`),GROMACS v2023.1 (`GRO`), MDAnalysis v2.1.0 (`MDA`) and LiPyphilic v0.11.0 (`LiP`) were used in this study. All analyses were performed on a workstation equipped with an Asus Z10PE-D8 WS motherboard, two Intel Xeon E5-2698 v4 CPUs (each with 20 cores, 3.6 GHz), two NVIDIA GeForce RTX 4070 GPUs, and 128GB of DDR4 RAM, running Ubuntu 22.04.4.

### Metrics and analysis details

#### Trajectory processing

For all analyses, except for the calculation of the lateral diffusion coefficient, the raw trajectories were processed using a similar workflow. The aim was to generate clean trajectories centered on the membrane and free of periodic boundary condition (PBC) artifacts for subsequent analyses. The standard processing pipeline comprises two main stages. First, the GROMACS utility `gmx trjconv` was used to extract the membrane-only trajectory, thereby removing all water and ion atoms^11,12^. Second, the resulting membrane-only trajectory was processed using CPPTRAJ^9^ in a three-step sequential pipeline:

1. `unwrap *`: All lipid molecules were first made whole, correcting for any splits across PBC boundaries.
2. `center * mass origin`: The center of mass of the entire (now whole) membrane was translated to the coordinate origin.
3. `image origin center`: All molecules were “folded” back into the primary simulation box, ensuring a single, compact membrane centered at the origin.

The trajectories used for the analysis of the lateral diffusion coefficient via the mean squared displacement (MSD) were different. `CPP` and `GRO` processed the raw, unaltered trajectories directly, while the `MDA` and `LiP` workflows required a separate trajectory pre-processed using `CPP`’s `unwrap` command (as detailed below).

#### Area per lipid and bilayer thickness

The area per lipid (APL) represents the average cross-sectional area occupied by a single lipid molecule in the membrane. `CPP`, `GRO MDA`and `LiP`, adopt the same mathematical definition for APL, given by

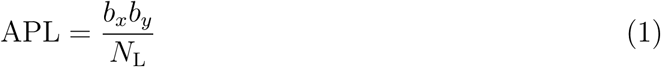

where *b_x_* and *b_y_*are the box dimensions along the *x* and *y* axes, respectively, and *N*_L_ denotes the number of lipids per leaflet. The values of *b_x_* and *b_y_* were extracted directly from the trajectory for each frame, using the `vector box` command in the case of `CPP`, the `gmx traj -ob` command in `GRO`, and the `Universe.dimensions` function in `MDA`, and `LiP`. *N*_L_ was calculated by dividing the total number of lipids—specified by the user—by two, assuming a symmetric leaflet distribution^9^.

The membrane thickness is typically defined as the distance between the head groups of opposing leaflets. For this metric, in `CPP` we first extracted the coordinates of all phosphorus atoms using the `trajout` command. A subsequent `awk` script implemented a dynamic leaflet assignment by calculating, for each frame, the center of mass (COM) along the *z*-axis (*z*-COM) for all phosphorus atoms. This *z*-COM was then used as a dividing plane to dynamically assign atoms to the upper and lower leaflets. The thickness was computed as the absolute distance between the *z*-COM of the upper leaflet atoms and the *z*-COM of the lower-leaflet atoms. An analogous “split-by-mean” dynamic leaflet assignment was implemented in `MDA`, in this case using the `center_of_mass()` function together with additional Python routines. `GRO` calculated thickness as the distance between the COMs of two static, pre-defined index groups representing the phosphorus atoms of the upper (`P_Upper`) and lower (`P_Lower`) leaflets. The *z*-component of the COM for each group was calculated for each frame using the `gmx traj -com` command^11,12^.

Finally, for thickness analysis in `LiP`, the `AssignLeaflets` class from the lipyphilic. leaflets.assign_leaflets module was used^15^. The class was instantiated once on the entire pre-processed trajectory using a selection of all lipid residues. The `assigner.run()` method was then called to generate a frame-by-frame record of the leaflet assignments for each lipid. Once these assignments were established, the thickness was computed for each frame. The analysis script iterated through the trajectory, retrieved the leaflet assignments for the current frame, grouped the lipids into upper and lower leaflets, and selected phosphorus atoms for each leaflet. The thickness was then calculated as the absolute distance between the *z*-center of mass of the upper leaflet phosphorus atoms and the *z*-center of mass of the lower-leaflet phosphorus atoms.

#### Deuterium order parameter

The deuterium order parameter for each lipid tail chain, |*S*_CD_|, is defined as

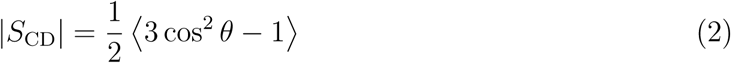

where *θ* is the angle between the C – H bond vector and the bilayer normal^26,27^. Although Equation 2 does not always reproduce the experimental values quantitatively, it is widely used to assess the differences in lipid tail disorder across systems and conditions^28,29^.

In `CPP`, |*S*_CD_| was calculated using a command (`lipidscd`) specifically designed to evaluate the order parameters of lipid bilayers^9^. `GRO` performed equivalent calculations (via the `gmx order` command).

For `MDA`, a custom script was developed to calculate |*S*_CD_|. The script first identified all C – H pairs by locating the carbon atoms in the sn-1 and sn-2 acyl chains and determining the bonded hydrogen atoms. It then iterated over each trajectory frame to compute the vector for every C – H pair and calculated the angle *θ* between each vector and the bilayer normal. The order parameter was then obtained by applying Equation 2, and the reported |*S*_CD_| values correspond to the time-averaged results over the entire trajectory.

`LiP`, in contrast, provides only a coarse-grained order parameter for each acyl chain (computed using the `lipyphilic.analysis.order_parameter` module^15^), thus reflecting the orientation of entire chain segments instead of individual carbon atoms along the chain. Consequently, these values are not directly comparable to atomistic calculations, and the results from `LiP` were not included in this part of the study.

#### Lipid headgroup orientation

The orientation of the lipid headgroup defines the interface between the hydrophobic core of the membrane and the surrounding aqueous environment. It is quantified as the angle between the headgroup vector—spanned by the phosphorus (`P31`) and nitrogen (`N31`) atoms—and the normal to the bilayer surface. The closer this angle is to 90°, the more parallel the vector lies relative to the membrane plane^30^.

The headgroup orientation was computed individually for each lipid by calculating the angle between the P– N vector (defined from the positions of `P31` and `N31`) and the bilayer normal (*z*-axis). In `CPP` this was achieved using the `vectormath` command with the `dotangle` function;^9^ in `GRO` the dedicated `gmx gangle` tool was used;^11,12^ and in the case of `MDA` the `MDA Universe` reader was employed in conjunction with NumPy functions as part of a custom Python script. `LiP` performed equivalent calculations using its dedicated `ZAngles` analysis class (from the `lipyphilic.analysis.z_angles` module^15^), initialized with the same `P31` and `N31` atom selections.

#### Lateral Diffusion coefficient

The lateral diffusion coefficient of lipids in the plane of the membrane, *D*_L_, can be calculated using the Einstein relation,

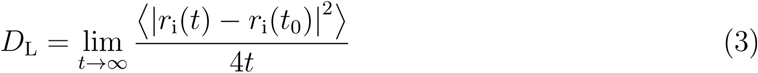

where *r*_i_(*t*) and *r*_i_(*t*_0_) represent the positions of atom *i* at times *t* and 0, respectively, and 〈|*r*_i_(*t*) − *r*_i_(*t*_0_)|^2^〉 is the MSD in the *xy* plane for the time *t* starting from *t*_0_ ^8^.

`CPP` used the dedicated `diffusion` command, which, when applied to the raw trajectory, internally handled molecule unwrapping and COM drift correction. The MSD for phosphorus atoms was then calculated separately in the *x* and *y* directions. The *x* and *y* components were summed to obtain the two-dimensional lateral MSD as a function of time, which was then saved for further analysis^9^.

`GRO` also employed a native tool `gmx msd` that internally accounts for motion across periodic boundaries^11,12^. In this workflow, the membrane subsystem was first extracted from the raw trajectory, and `gmx msd` was applied to the membrane-only trajectory.

The workflows for `MDA` and `LiP` were found to be identical. This analysis uniquely required a separate preprocessing step: the raw, unprocessed trajectories were first treated with `CPP` using its `strip` and `unwrap` commands to create a temporary, unwrapped, membrane-only trajectory file. This file was then read by the Python scripts, and the MSD was calculated using the `MDAnalysis.analysis.msd.EinsteinMSD` class applied to the phosphorus atoms over time. This was done with the `select_com=None` parameter, which disabled the internal COM-drift subtraction of the class to mimic the behavior of the `CPP diffusion` command.

After obtaining the MSD, a custom script performed a least-squares fit on the linear portion of the MSD versus time curve. The fitting interval spanned from 10% to 50% of the total simulation time, excluding both the ballistic regime at short time lags and the poorly averaged data at longer time scales. The slope of the fitted line was then used to calculate *D*_L_ according to Equation 3^31^. This same fitting procedure was applied in post-processing to the raw MSD data generated by `CPP` and `GRO`.

### Simulation Convergence and Inter-Replicate Equivalence

To validate the simulation data, we performed a two-part analysis assessing convergence and inter-replicate equivalence on the 500 ns time series (2500 frames), using the CPP data for APL and bilayer thickness as a representative set.

First, intra-replicate convergence was evaluated. The cumulative averages of both properties showed a stable plateau over the course of the simulation time (Figs. SI.1–SI.3). Furthermore, an analysis of 10 sequential 50 ns blocks showed that the block averages fluctuated consistently around the overall mean, and the autocorrelation function for each replica rapidly decayed to zero. Therefore, 500 ns were considered sufficient for parameter convergence, and the full simulation time was suitable for parameter calculation.

Second, the inter-replicate statistical equivalence was assessed. A comparison of the block averages over time confirmed that all three replicates evolved similarly and sampled consistent and equivalent thermodynamic states. The data from the three replicates were thus considered sufficiently converged to be pooled for subsequent software analysis and comparison, which was based on a frame-by-frame average across the three replicates.

## Results and Discussion

### Influence of membrane size and hydration

The influence of membrane size and hydration level on bilayer properties was investigated. Simulations on 64L–*j*W and 128L–*j*W with *j* = 40, 80, 160 systems were carried out but excluded from the final analysis due to severe finite-size artifacts. The 64L systems failed to maintain bilayer structural integrity, likely as a result of self-interaction artifacts arising when the box dimensions approached the nonbonded cutoff radius. The 128L systems exhibited pronounced anisotropic deformation, drifting from the initial square geometry to highly elongated rectangular box shapes. Consequently, the 256L system was determined to be the minimum size required to ensure stable and geometrically realistic trajectories.

The results summarized in Figure 1a suggest that the APL values for different hydration levels (40W, 80W, and 160W) are not statistically different. The group medians were very close, the boxes overlapped almost completely across all hydration levels, and the variability within each group was larger than the variation between different hydration conditions. These findings differ from those of previous studies. Castro-Román et al. ^4^ reported that APL is influenced by hydration for DOPC bilayers, Farhangian et al. ^7^ showed that APL increases with hydration for DOPE, POPE, DOPC and POPC. However, in all these studies, the hydration levels were significantly lower—typically below 30W—than those present in our work—40W and above. Taken together, these observations suggest that, at least for POPC membranes, hydration levels near 40W mark the transition to the fully hydrated state, where the increase of hydration no longer affects the bilayer’s structural properties within the same membrane size.

**Figure 1.**
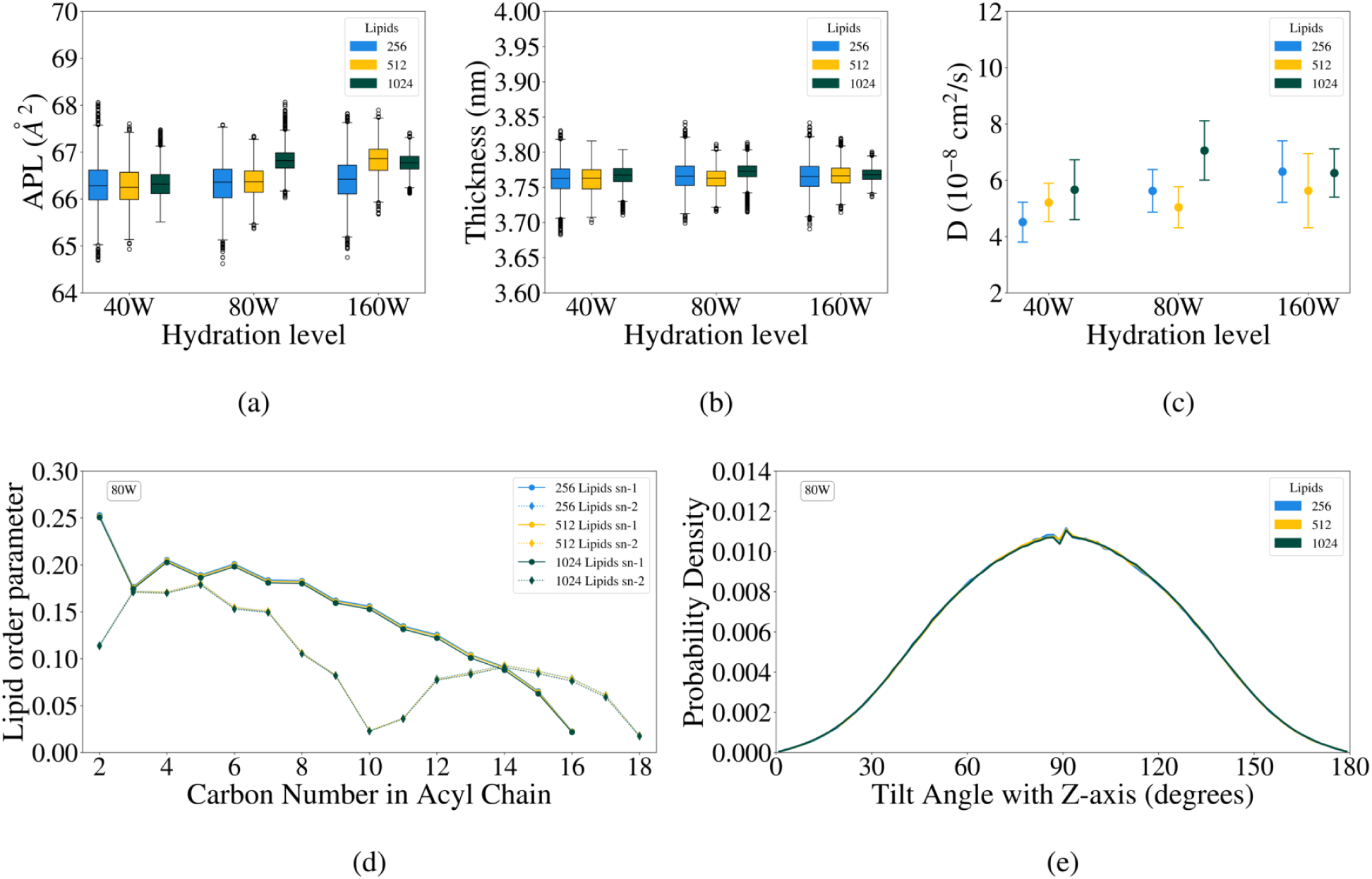
Influence of membrane size and hydration level on bilayer properties. **(a)** Area per lipid (APL) and **(b)** bilayer thickness show a consistent decrease in data variance (interquartile range) as membrane size increases, indicating greater statistical stability in larger systems. **(c)** Lateral diffusion coefficients display a size-dependent increase specifically at the 40W hydration level, whereas higher hydration levels show no pronounced correlation. **(d)** Lipid order parameters and **(e)** headgroup orientation (80W systems shown as representative) exhibit characteristic profiles with no measurable influence from bilayer size.

The results for the membrane thickness (Figure 1-(b)) also shows that both the hydration level and number of lipids yielded no statistically significant differences between the two groups. However, the larger the system, the greater the precision in both cases, that is, a trend that is consistent with the established finite-size effects in membrane simulations, where the probability distributions of structural properties are known to depend on the system size^26^.

The lateral diffusion coefficient (Fig. 1-(c); see Fig. SI.4 to SI.6 for the corresponding MSD plots) increases with hydration and exhibits small but visible differences between the 256L system and the larger systems, consistent with the finite-size effects that are expected to affect dynamic properties more strongly. This suggests that the hydration level is a determining factor in whether and how the system size affects lipid dynamics. While Klauda et al.^5^ reported a decrease in diffusion with increasing system size in fully hydrated bilayers due to short-range collisional correlations, our results at lower hydration (40W) show the opposite trend. The observed increase in lateral diffusion with bilayer size at 40W is consistent with the hydrodynamic finite-size scaling effects described by Vögele et al.^32^, in which PBC suppress diffusion in smaller systems through solvent-mediated interactions. Additionally, larger membrane patches can accommodate long-range phenomena, such as membrane undulations and mesoscopic hydrodynamic fluctuations, which are suppressed in smaller systems as detailed by Ayton and Voth^33^.

Figure 1 also includes the results for the acyl chain order parameter and tilt angle distribution. The order parameter profiles for the three system sizes overlapped almost perfectly, indicating good convergence of the conformational properties across system sizes. A typical trend was observed, that is, a plateau in the initial carbons (4–6), followed by a decrease towards the chain end^34^. No changes were observed across the hydration levels, which is expected for fully hydrated bilayers, where the hydrophobic core is largely insensitive to the amount of water in the bulk. The lipid tilt-angle distributions were also similar.

### Software comparison and performance

The 1024L–80W system was selected as a representative case for comparing the outcomes produced by the different software packages. The results are summarized in Figure 2 (see Figs. SI.7–SI.9 for all *i*L–*j*W systems).

**Figure 2.**
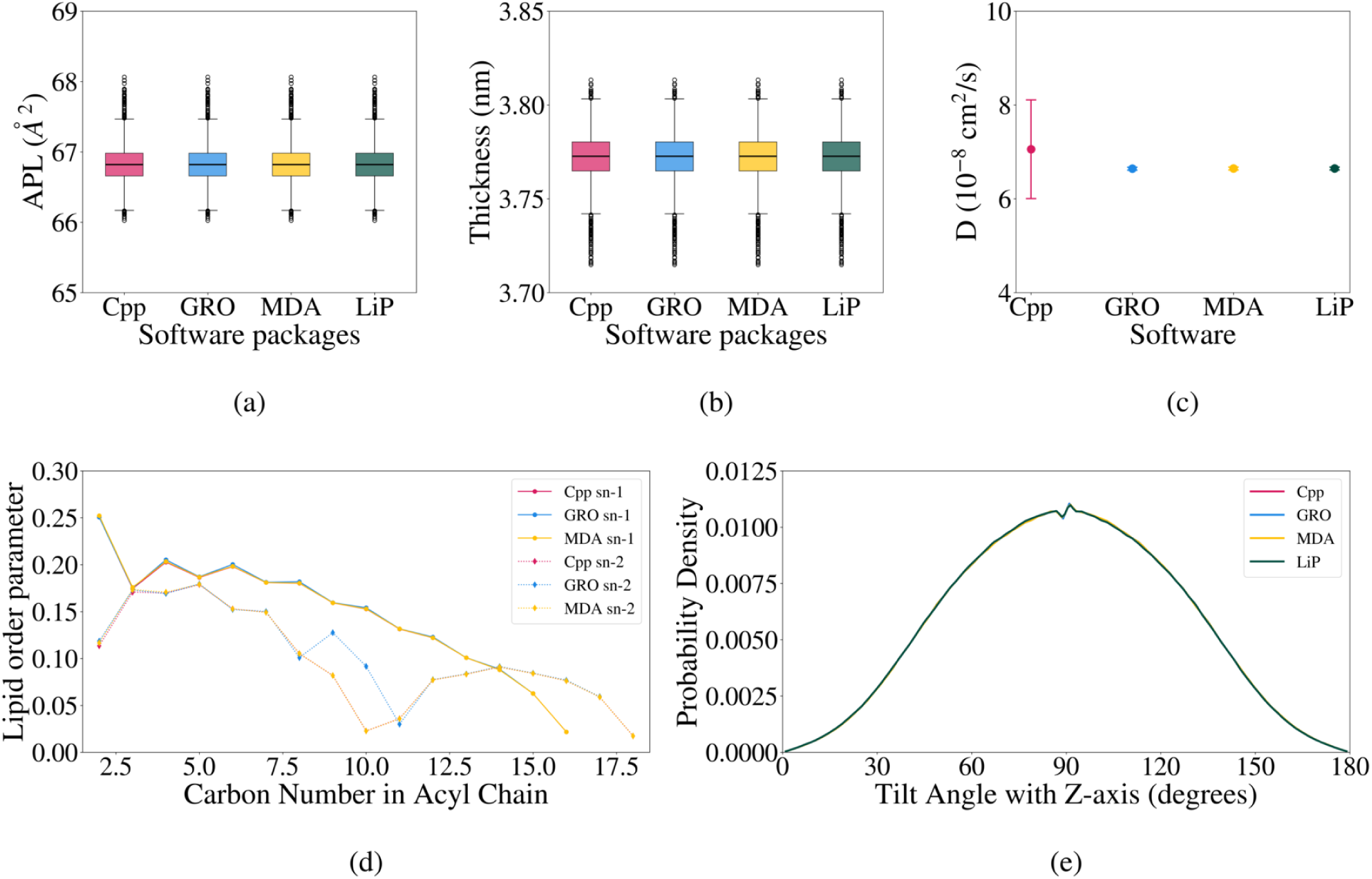
Comparative analysis of bilayer properties for the 1024L–80W system across different software packages. **(a)** APL and **(b)** bilayer thickness show no statistical or visual differences between tools. **(c)** Lateral diffusion coefficients exhibit strong agreement among `GRO`, `MDA`, and `LiP`, whereas `CPP` shows a larger standard deviation due to the absence of time-origin smoothing. **(d)** Lipid order parameters are consistent for saturated carbons, but `GRO` produces an artifact at the sn-2 double bond (C9–C10) due to its known limitation with unsaturated lipids (`LiP` excluded). **(e)** Lipid headgroup orientation is identical across packages.

For APL, the analysis revealed no significant differences between the software packages. All tools implemented the same mathematical expression (Eq. 1) and, as expected, produced consistent APL averages with nearly indistinguishable distributions. However, it is important to note that this standard geometric approach treats the membrane as a flat surface, thereby disregarding the local curvature inherent to dynamic bilayers. To complement this, a Voronoi tessellation analysis accounting for 3D surface curvature was performed on the 1024L–80W system, averaging the results of the last frame across three replicates (see Fig. SI.10). This method explicitly accounts for *z*-axis undulations and yielded a larger average APL (67.9 Å^2^) compared to the geometric projection (66.6 Å^2^). The difference of approximately 1.36 Å^2^ indicates that the geometric approach in Equation 1 only slightly underestimates (∼2%) the true contact surface area available for solvent and lipid interactions in large, undulating membranes. Both the standard geometric and the Voronoi tessellation approaches produced APL values slightly higher than the experimental APL of 63.3 Å^2^ reported by Kučerka et al.^35^. The latter value was derived using the Scattering Density Profile (SDP) model, which is not a direct measurement but rather a method developed to reconcile discrepancies in APL obtained from standalone X-ray and neutron scattering experiments^36^.

The boxplots of membrane thickness were also visually indistinguishable, indicating that for a stable, well-equilibrated bilayer, all four calculation methods produced identical results. A notable exception was observed for the 1024L–40W system (Fig. SI.9), which exhibited high-value outliers in the `MDA` results. Given the large membrane size combined with the lowest hydration level—conditions that promote stronger membrane undulations—these outliers suggest a limitation of the frame-by-frame geometric “split-by-mean” algorithm in correctly partitioning the bent membrane. The average thickness of 3.77 nm returned by all packages is slightly lower than, but comparable to, the experimental value of 3.91 nm reported by Kučerka et al.^35^. This difference is consistent with the larger simulated APL relative to the SDP model discussed above, in which the membrane thickness is proportionally reduced to conserve lipid volume.

The order parameter profiles for the sn-1 chain showed a high degree of consistency between the `GRO`, `MDA`, and `CPP`simulations. All three tools captured the characteristic plateau for the upper carbons (4-6) and the progressive decrease in order towards the methyl end. However, a significant discrepancy appeared in the sn-2 acid chain, which contained a *cis*-double bond at the C9-C10 position. `GRO` produced an artificial increase at these carbons. This is a known artifact of the `gmx order` tool, which is designed for saturated carbons and relies on a static index file, making it unsuitable for unsaturated lipids^37^. The tool documentation explicitly states that it only works for saturated carbons ^11^. This limitation also explains why the `GRO` sn-2 chain calculation terminated at C10 and why the sn-1 chain calculation omitted the data for the terminal methyl group. To address these and other limitations of `gmx order`, alternative and more robust GROMACS-compatible tools such as `gorder` have been developed^37^.

For the lateral diffusion coefficient of the 1024L–80W system, the tools showed good methodological agreement. `GRO`, `MDA`, and `LiP` produced nearly identical diffusion coefficients with low standard deviation across the three replicates. In contrast, the native `CPP diffusion` command yielded a larger coefficient with a noticeably higher standard deviation, indicating lower *inter* -replicate consistency. This discrepancy arises from differences in the statistical averaging protocols. `GRO`, `MDA`, and `LiP` employ time-origin shifting to compute the MSD^38^. By averaging over multiple time origins, these algorithms accelerate the statistical *intra*-replicate convergence of the diffusion coefficients. This internal averaging drives the results of physically equivalent replicates toward the ensemble mean, producing the low *inter* -replicate standard deviations observed. Conversely, `CPP` calculates displacements directly from the raw coordinates, without an implicit time-averaging layer. As a result, the `CPP` coefficients are less statistically converged between replicates, and the corresponding MSD curves appear less smooth than those produced by the other packages (Figs. SI.4–SI.6). However, this trend was reversed for systems at the extremes of bilayer size and hydration—namely 256L—160W and 512L–40W—where `GRO`, `MDA`, and `LiP` exhibited higher standard devia-tions than `CPP` (Figs. SI.7–SI.9). In these cases, the *intra*-replicate smoothing algorithms in `GRO`, `MDA`, and `LiP` filtered out stochastic noise, revealing genuine *inter* -replicate heterogeneity, whereas the noise inherent to the raw `CPP` analysis partially masked these differences. These results are consistent with the differences in the analysis workflows: as detailed in the Methods, `CPP` and `GRO` processed the raw trajectories directly using their internal unwrap-ping logic, whereas the `MDA` and `LiP` workflows—functionally identical to each other—used trajectories pre-processed with the `CPP unwrap` command. This demonstrates that the `MDA EinsteinMSD` class (with `select_com=None`) produced results nearly identical to those from `GRO`’s `gmx msd` but distinct from those of `CPP`’s `diffusion` command, even when operating on a `CPP`-generated input trajectory. Consequently, the standard deviation reported in the lateral diffusion coefficient subplot of Figure 1 provides a transparent representation of the system’s stochastic variability, capturing genuine sampling differences between independent replicates rather than artifacts introduced by smoothing algorithms.

An important yet subtle feature of the analysis packages was uncovered during the MSD calculations. In preliminary tests, corrupted trajectories (containing inconsistent time meta-data) were inadvertently used as inputs for the lateral diffusion analysis. The `GRO` tool (`gmx msd`) was the only routine that failed and returned an error indicating the corrupted time data. This behavior arises because `gmx msd` natively relies on the trajectory’s internal frame/time metadata and checks its consistency. In contrast, the `CPP diffusion` command and the `MDA`-based script bypass the internal metadata by applying a user-specified time step to the sequence of frames. As a result, they can—and do—proceed to produce erroneous outputs without issuing any warning. The `gmx check` utility was therefore essential for diagnosing inconsistencies in trajectory time metadata^11^. All results presented in this paper for all analysis tools are based on trajectories that were verified to be consistent and non-corrupted.

To assess computational performance, the 1024L–80W system was benchmarked using all four software packages. The tests were executed with parallel jobs pinned to specific physical cores, ranging from 1 to 20 cores, on a single NUMA node to ensure consistent measurements^39^. Because the `CPP` and `GRO` tools used for these analyses are serial, they were benchmarked only on a single core^11,21^. The Python-based `MDA` and `LiP` scripts were adapted to use Python’s `multiprocessing` library to parallelize the calculations^15,40^. Figure 3 shows the resulting CPU time as a function of the number of cores.

**Figure 3.**
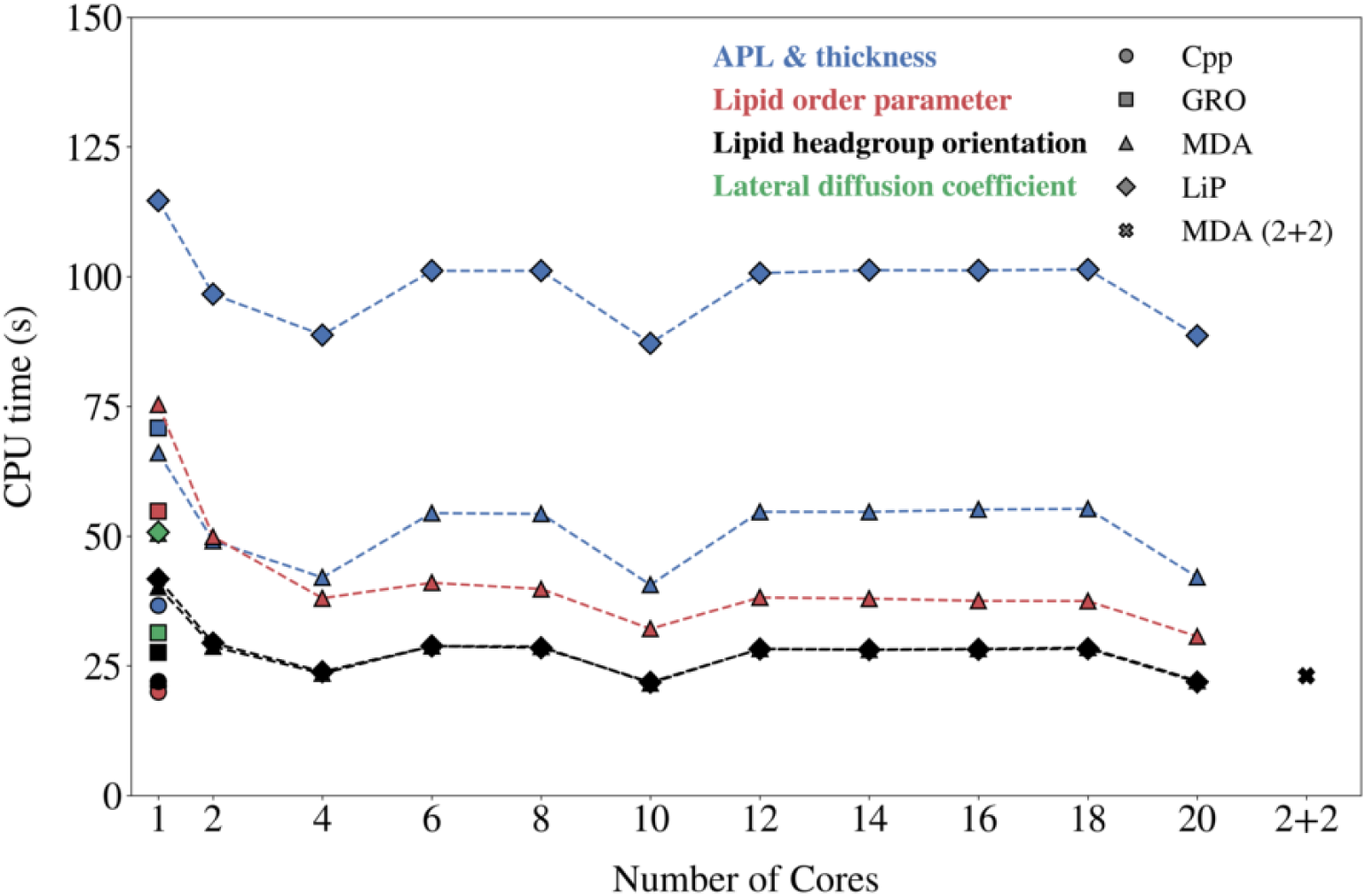
Computational performance benchmark showing CPU wall-clock times for analyzing the 1024L–80W system. `CPP` consistently exhibits the fastest single-core performance across all metrics. Python-based tools (`MDA` and `LiP`) show substantial speedups with parallelization (up to 20 cores), although `LiP`’s scaling for thickness calculations is limited by a fixed serial topology-scanning step. Lateral diffusion calculations were restricted to single-core execution for all packages.

For the combined APL and bilayer-thickness performance, `CPP` was the fastest serial tool, followed by `MDA` and `GRO`, which showed comparable run times, whereas `LiP` was significantly slower than the others. Parallel execution introduced a substantial, though nonlinear, advantage for both `MDA` and `LiP`. For the APL calculations, the parallel `MDA` and `LiP` scripts were functionally identical; the marked difference in total runtime arose from the bilayer-thickness calculations. The `MDA` script implemented the “split-by-mean” geometric method, which was fully parallelizable because each frame could be processed independently. In contrast, the `LiP` script uses the topological `AssignLeaflets` routine, which must first be executed serially to scan the entire trajectory and construct the leaflet-assignment map. Subsequent frame-wise thickness calculations were parallelized using this precomputed map. Consequently, the `LiP` performance profile reflected a scaling benefit that was partially offset by a fixed serial overhead from the `AssignLeaflets` step—an overhead absent from the fully parallel `MDA` workflow. Furthermore, the best performance achieved by `MDA` was observed at 10 and 20 cores, comparable to that of `CPP` running on a single core. `LiP` exhibited the same qualitative scaling behavior as `MDA` despite being significantly slower across all core counts.

For the order-parameter analysis, `CPP` substantially outperformed both `GRO` and the `MDA`-based scripted approach on a single CPU core. The more computationally demanding `MDA` script benefited greatly from parallelization, achieving its best performance at 10 and 20 cores. This speedup was sufficient to outperform the serial `GRO` calculation but still fell short of matching the single-core performance of the `CPP lipidscd` command.

Lipid headgroup–orientation calculations were also fastest with `CPP` on a single core, followed by `GRO`, `MDA`, and `LiP`. Both the `LiP` and `MDA` scripts used equivalent custom-parallelized Python/NumPy functions and achieved their best performance at 10 and 20 cores, respectively, at which point their runtimes were indistinguishable from that of the serial `CPP` run. The special 2+2 core test—in which four cores are split evenly across two NUMA nodes—yielded a runtime indistinguishable from the 4-core single-node test, indicating that cross-node memory access is not a performance bottleneck.

Only single-core calculations were possible for the lateral diffusion coefficient across all software packages. `CPP` was again the fastest, followed by `GRO`. The `MDA` and `LiP` workflows were the slowest and, being functionally identical, exhibited indistinguishable run times.

## Conclusion

Our results show that the average values of APL, bilayer thickness, lipid-tail order parameters, and headgroup orientation were largely independent of membrane size and hydration over the ranges examined. The primary effect of increasing system size was a systematic reduction in the statistical variance of both APL and thickness, consistent with the expectation that larger systems yield more robust statistical averages by sampling a greater number of lipids. The lateral diffusion coefficient was also sensitive to membrane size and hydration, exhibiting a clear size dependence at 40W and a more complex, non-monotonic interplay between size and hydration at 80W and 160W. These variations are consistent with membrane undulations and hydrodynamic fluctuations that become more prominent as system size increases.

Across the four software packages, APL and membrane thickness were in excellent agreement, confirming that these equilibrium structural metrics are robust to implementation details when standard definitions are used. The most pronounced discrepancy arose in the order-parameter calculations, where the `GRO gmx order` tool produced a known artifact at unsaturated carbons, reflecting its documented inapplicability to unsaturated systems. Localized discrepancies were observed in one `MDA` thickness protocol for the 1024L–40W system and in the `CPP` diffusion calculation, underscoring the importance of understanding algorithmic assumptions when interpreting subtle differences between tools.

From a performance perspective, `CPP` was consistently the fastest serial tool across all five properties analyzed. Parallelization at 10 and 20 cores enabled the Python-based `MDA` and `LiP` workflows to approach—and in some cases match—`CPP`’s single-core performance for APL, thickness, and headgroup orientation, although `CPP` remained substantially faster for the order-parameter and lateral-diffusion calculations.

Taken together, these results provide practical guidance for simulation design and analysis. For studies primarily concerned with average structural properties, moderately sized systems (e.g., 256L), combined with a performance-optimized analysis package such as `CPP`, offer an efficient workflow that yields results comparable to those obtained from larger and more computationally demanding simulations. However, for dynamic properties, both system setup and the choice of analysis algorithms require additional scrutiny.

## Declaration of generative AI and AI-assisted technologies in the writing process

The authors used ChatGPT exclusively for language editing to improve the clarity and readability of the manuscript. Gemini 2.5 Pro was used to assist in the generation of Python and Bash scripts used for data analysis. All code was verified and validated by the authors. No generative AI tools were used for data analysis, interpretation, or drafting of scientific content. The authors remain fully responsible for all content and conclusions presented.

## Data Availability

Simulation trajectories (water removed, sampled at 1 ns/frame), provided in both continuous (for diffusion analysis) and centered/imaged formats, are available at DOI: 10.5281/zenodo. 17867493. This repository also contains all scripts used for parameter analysis with an accompanying guide, as well as the input and topology files necessary to reproduce the simulations.

## Acknowledgement

This work was completed in part with resources provided by the Auburn University Easley Cluster and University of Coimbra CERES research center, which is supported by FCT, within the projects DOI: 10.54499/UID/00102/2025 (Base and Programmatic funding) and DOI: 10.54499/UID/PRR/00102/2025.

## TOC Graphic

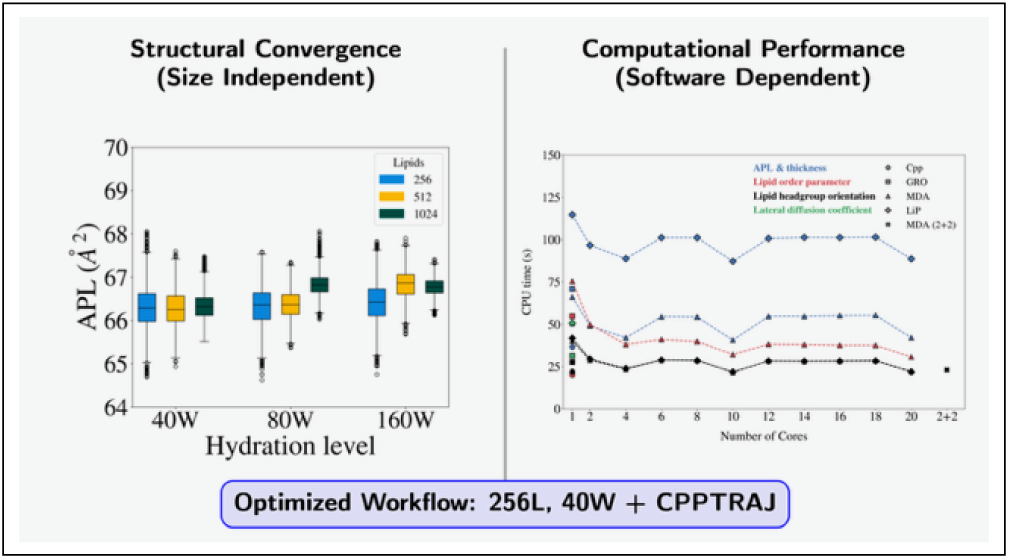

