## Supplementary material for "Bigger Isn’t Always Better: Comparing System Size, Hydration, and Software for Lipid Membrane Analysis": SI Figures 1-10 and SI Table 1

### Supporting Information

F. Carvalho,[†] P. Maximiano,[‡] P.N. Simões,[*,†] and M. Hashemi[*,‡]

†*University of Coimbra, CERES, Department of Chemical Engineering, Rua Sílvio de Lima, Coimbra, 3030-790, Portugal*

‡*Department of Physics, Auburn University, Leach Science Center 3126, Auburn, AL, 36849-5319, USA*

### Details of the simulated systems

Table SI.1: Initial box sizes and compositions of the simulation systems.

| System | Box size ($nm^3$) | POPC | $H_2O$ | $Na^+$ | $Cl^-$ | atoms |
|---|---|---|---|---|---|---|
| 128L-40W | $6.63 \times 6.62 \times 8.15$ | 128 | 5120 | 10 | 10 | 32532 |
| 128L-80W | $6.53 \times 6.56 \times 11.62$ | 128 | 10240 | 24 | 24 | 47920 |
| 128L-160W | $6.56 \times 6.54 \times 18.79$ | 128 | 20480 | 51 | 51 | 78694 |
| 256L-40W | $9.38 \times 9.41 \times 8.14$ | 256 | 10240 | 21 | 21 | 65066 |
| 256L-80W | $9.27 \times 9.24 \times 11.64$ | 256 | 20480 | 49 | 49 | 95842 |
| 256L-160W | $9.25 \times 9.26 \times 18.84$ | 256 | 40960 | 104 | 104 | 157392 |
| 512L-40W | $13.29 \times 13.31 \times 8.15$ | 512 | 20480 | 44 | 44 | 130136 |
| 512L-80W | $13.09 \times 13.08 \times 11.62$ | 512 | 40960 | 100 | 100 | 191688 |
| 512L-160W | $13.14 \times 13.09 \times 18.86$ | 512 | 81920 | 210 | 210 | 314788 |
| 1024L-40W | $18.83 \times 18.86 \times 8.17$ | 1024 | 40960 | 91 | 91 | 260278 |
| 1024L-80W | $18.51 \times 18.59 \times 11.65$ | 1024 | 81920 | 202 | 202 | 383380 |
| 1024L-160W | $18.57 \times 18.54 \times 18.87$ | 1024 | 163840 | 423 | 423 | 629582 |

### Convergence Analysis

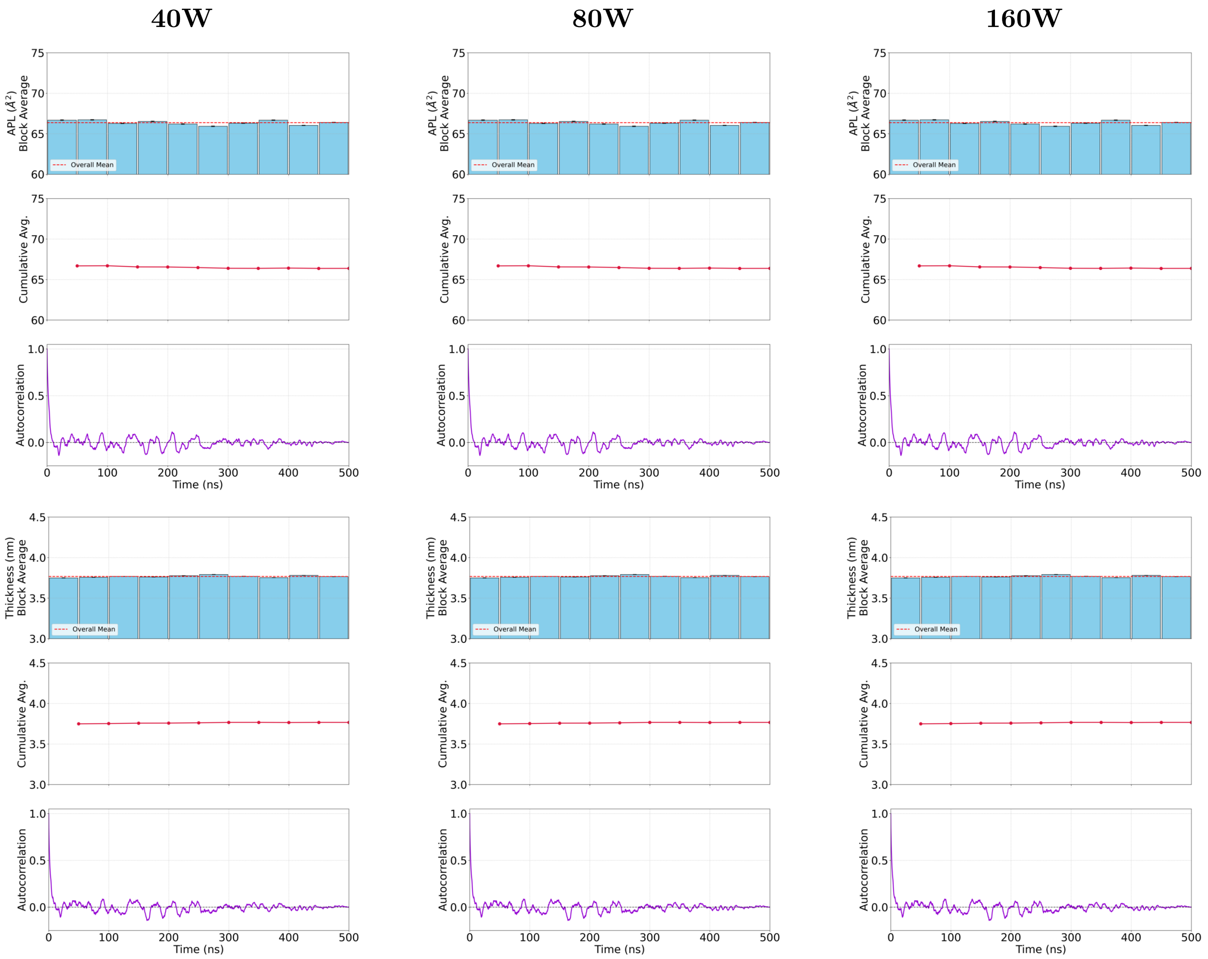


**(a)** Replica 1

**Figure SI.1.** Convergence analysis the 256L-$j$W systems. Columns correspond to hydration levels $j = 40, 80, 160$, from left to right. Plots show area per lipid (top three) and bilayer thickness (bottom three).

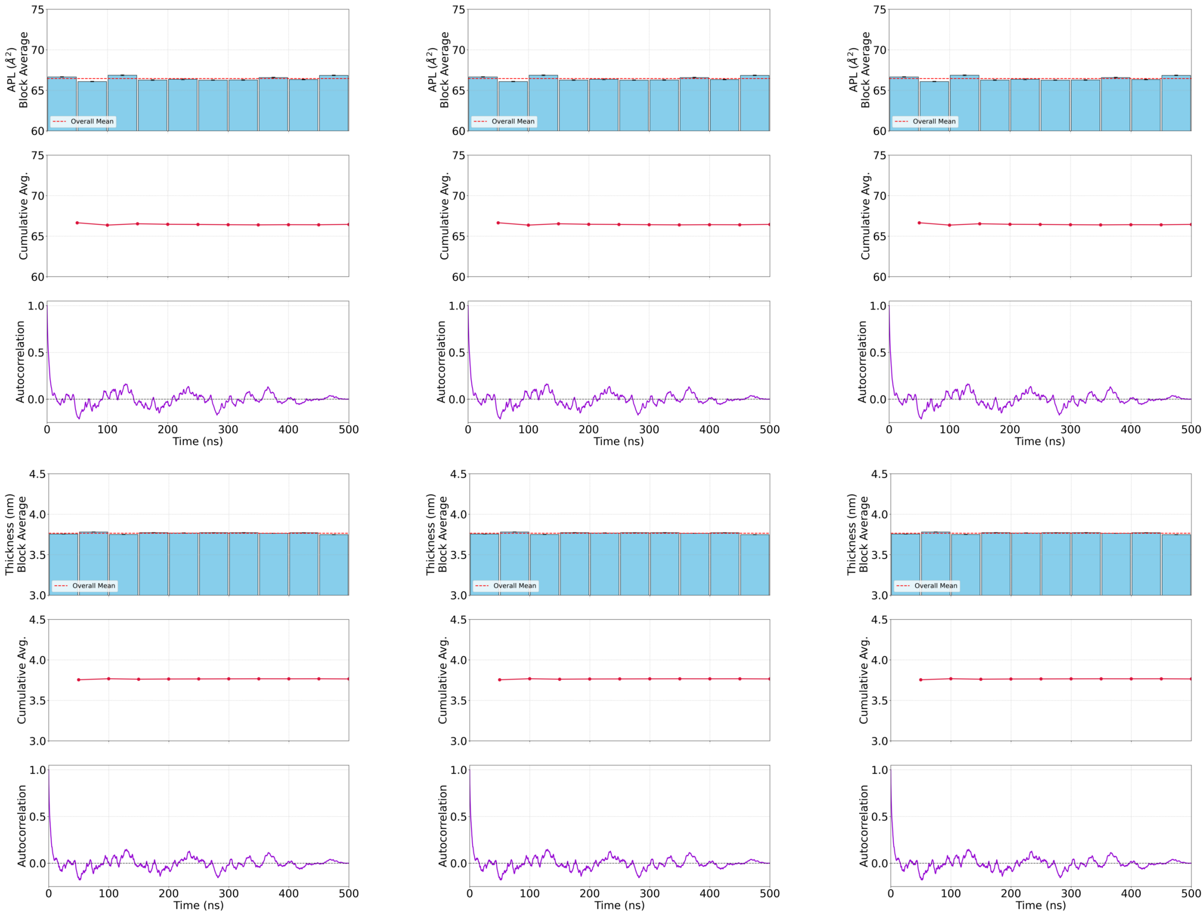


**(b)** Replica 2

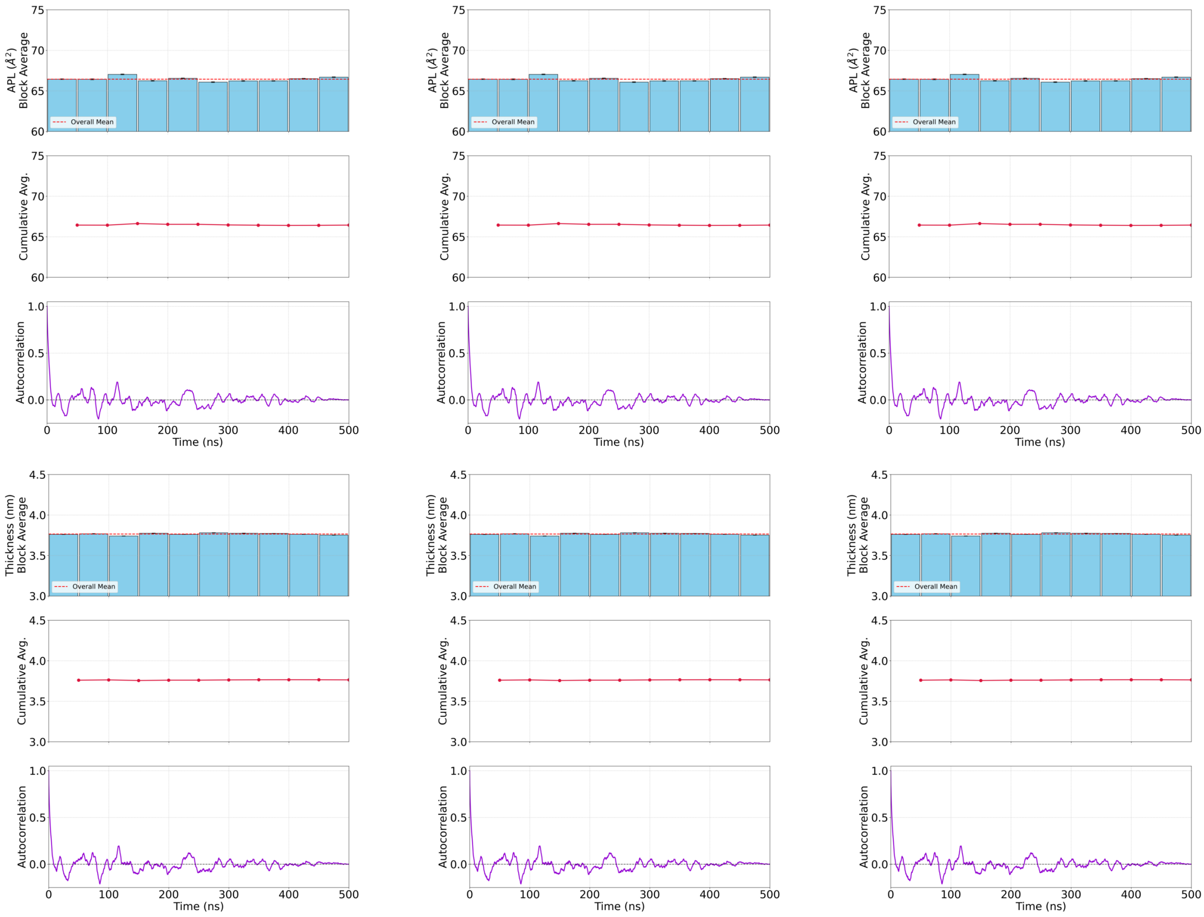


**(c)** Replica 3

**Figure SI.1.** Continued.

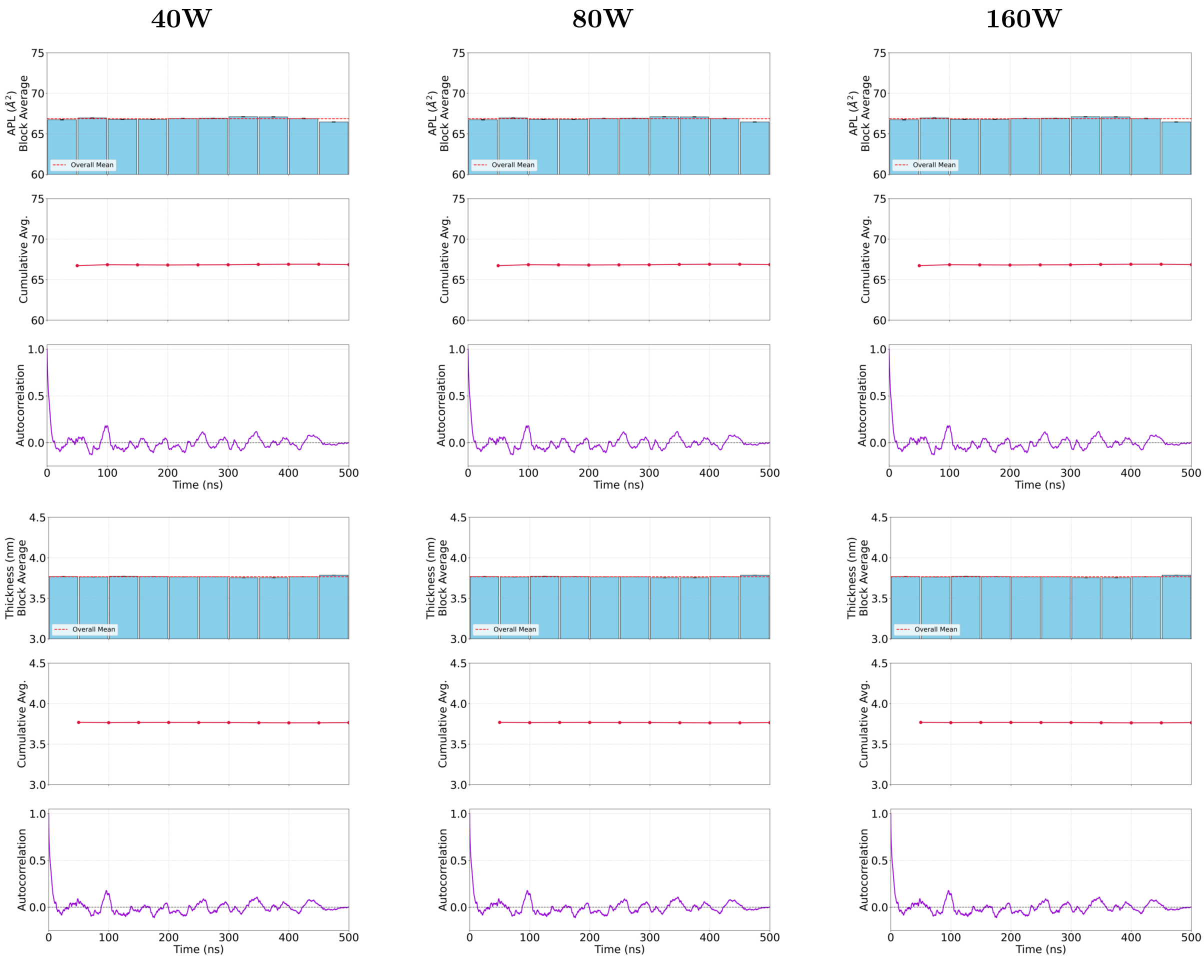


**(a)** Replica 1

**Figure SI.2.** Convergence analysis the 512L-$j$W systems. Columns correspond to hydration levels $j = 40, 80, 160$, from left to right. Plots show area per lipid (top three) and bilayer thickness (bottom three).

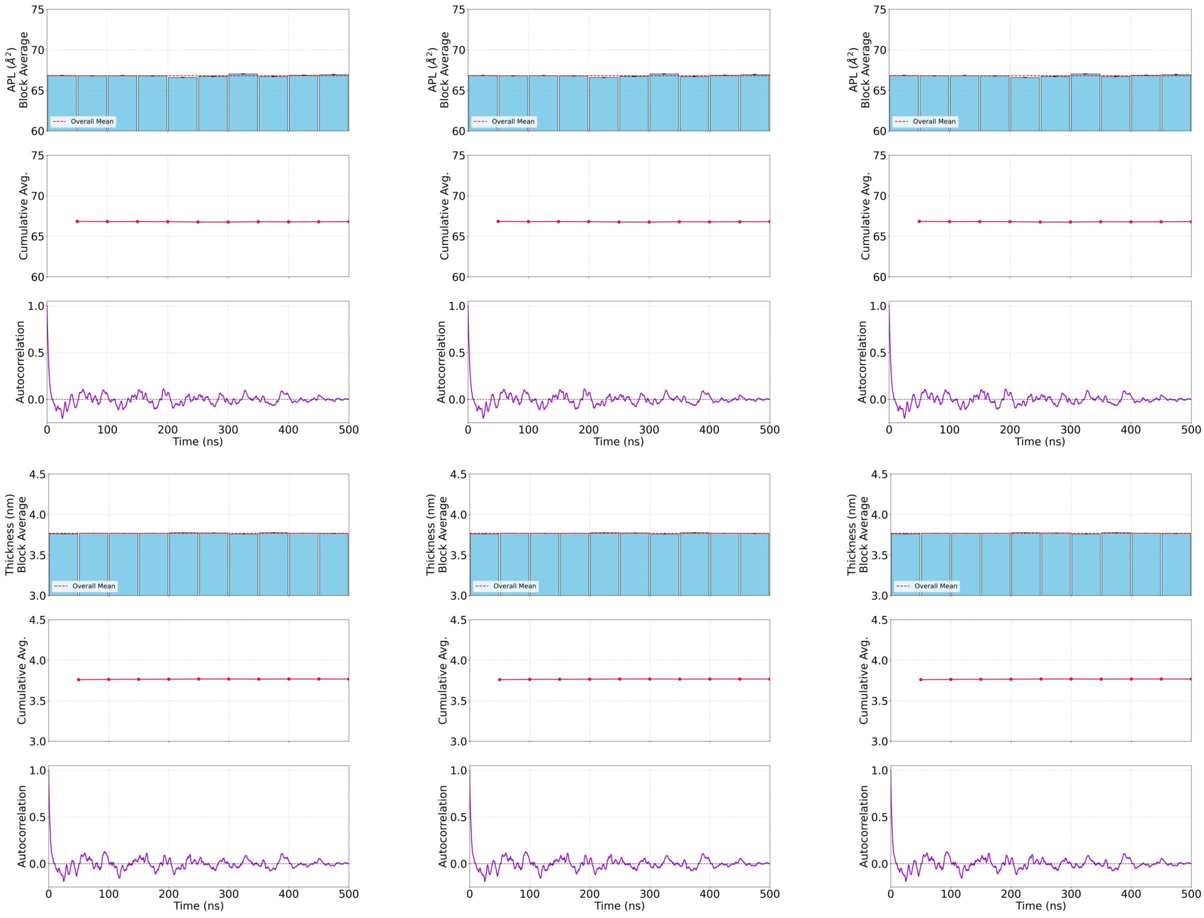


(b) Replica 2

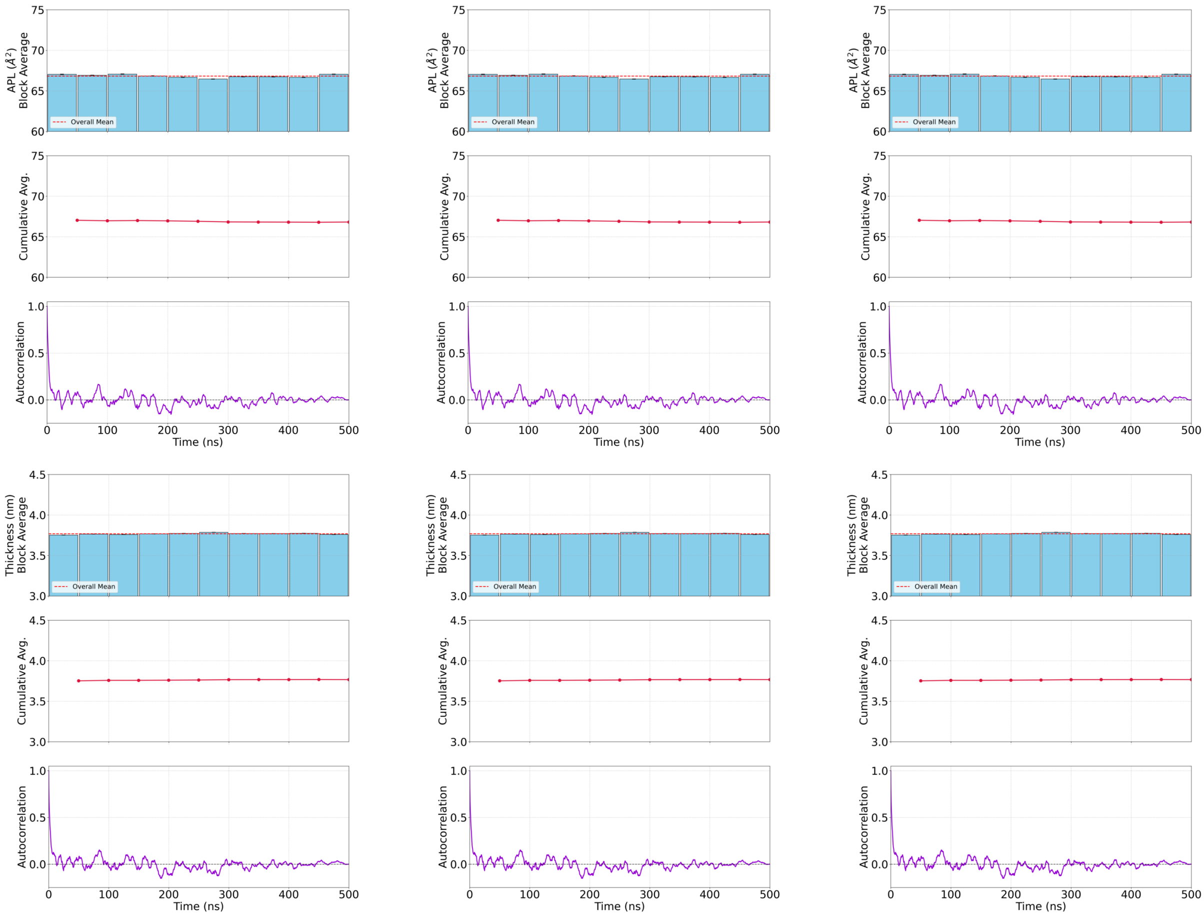


(c) Replica 3

**Figure SI.2.** Continued.

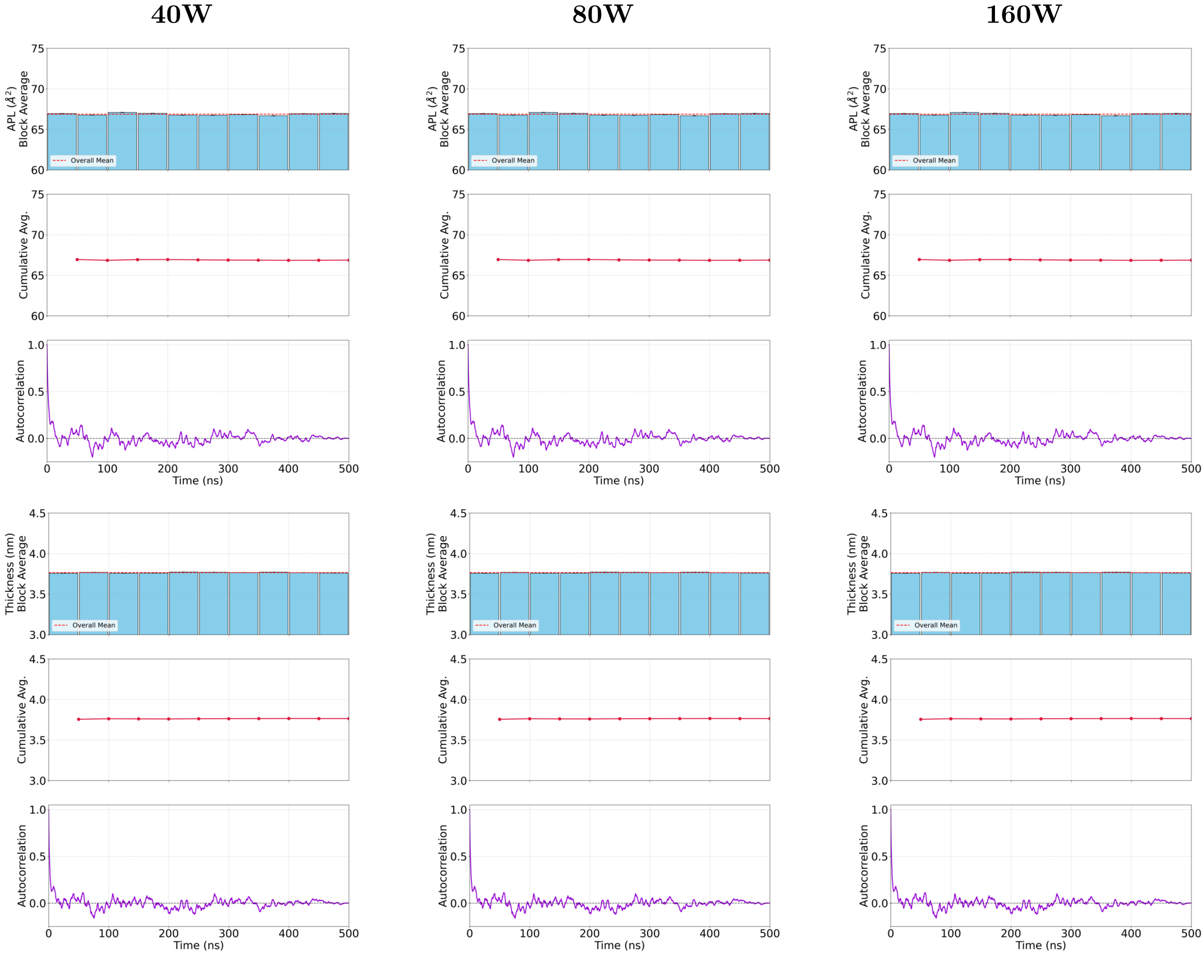


**(a)** Replica 1

**Figure SI.3.** Convergence analysis the 1024L-$j$W systems. Columns correspond to hydration levels $j = 40, 80, 160$, from left to right. Plots show area per lipid (top three) and bilayer thickness (bottom three).

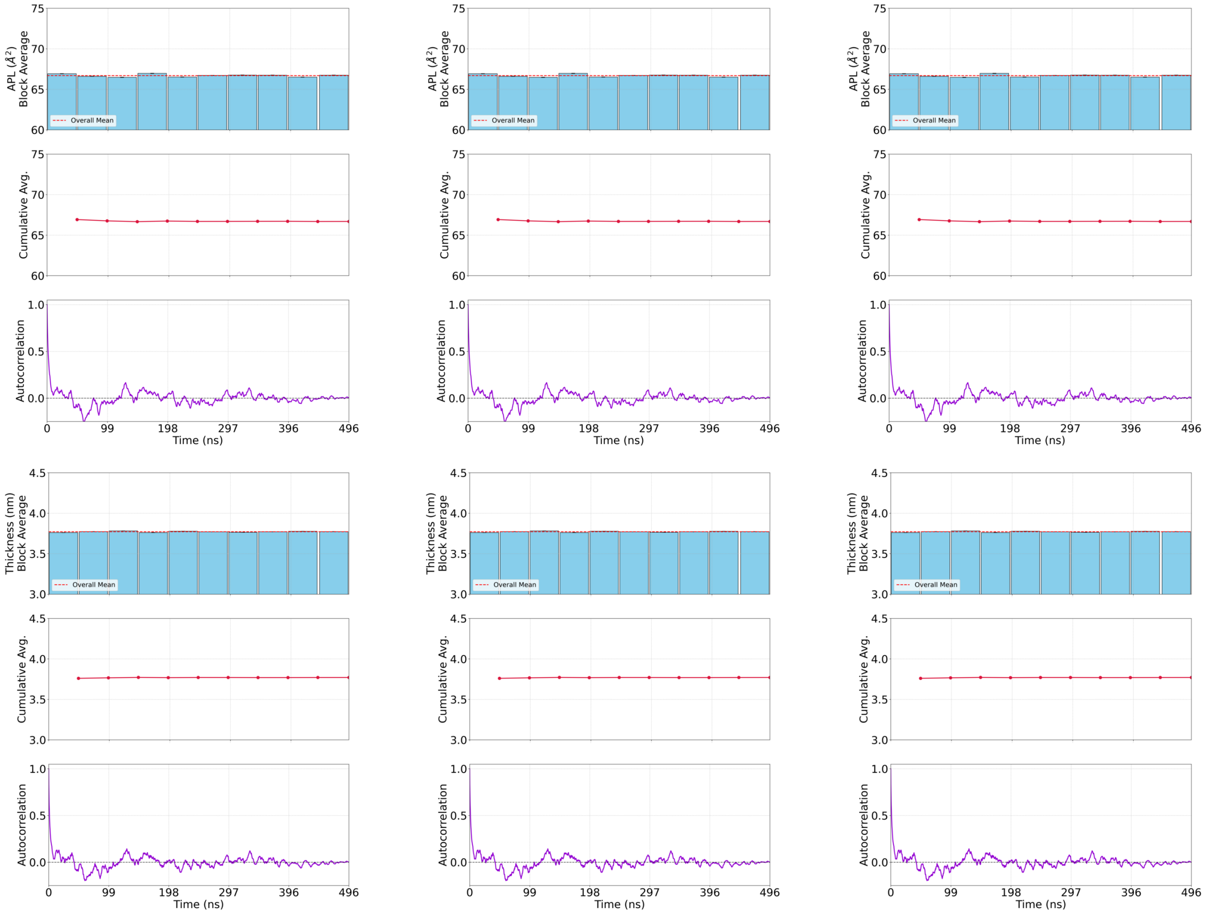


**(b)** Replica 2

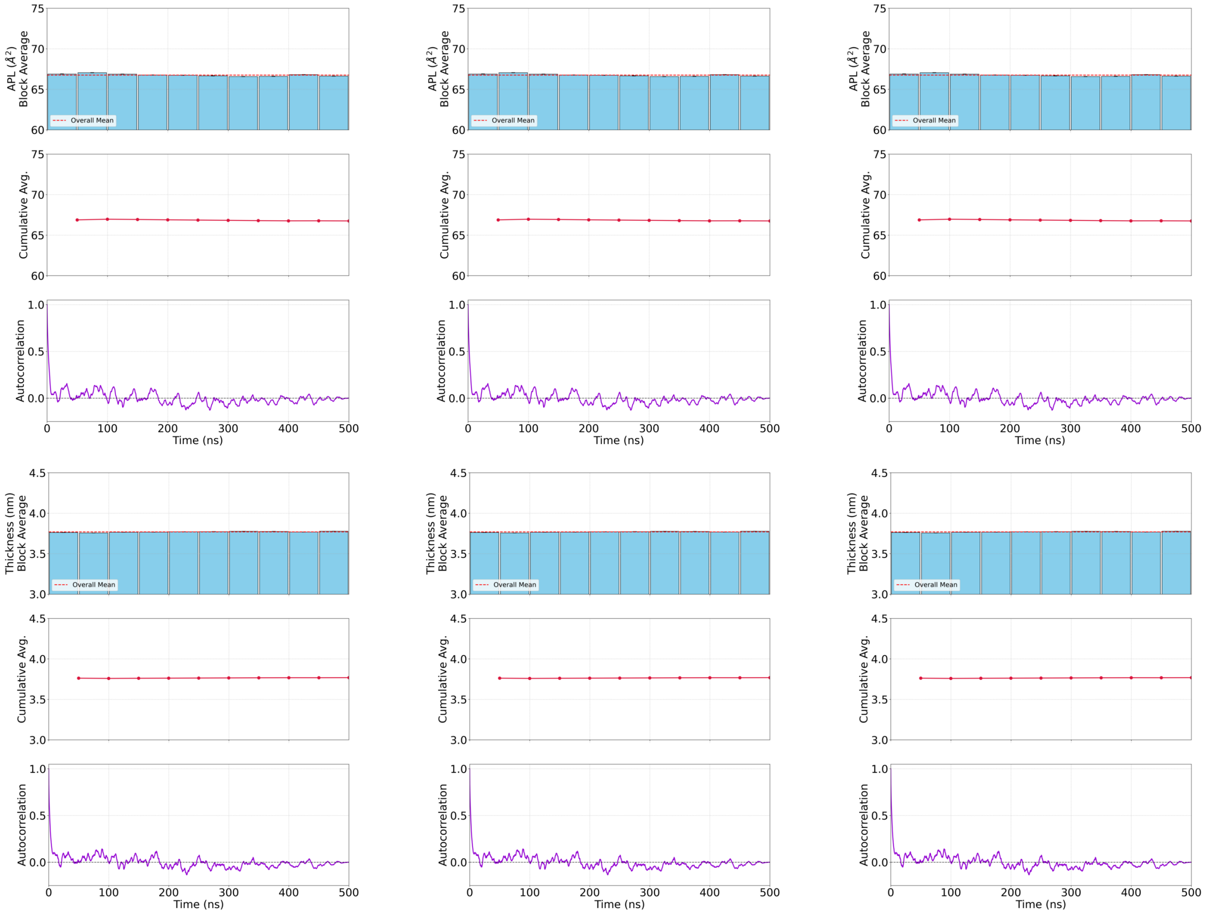


**(c)** Replica 3

**Figure SI.3.** Continued.

### MSD comparison

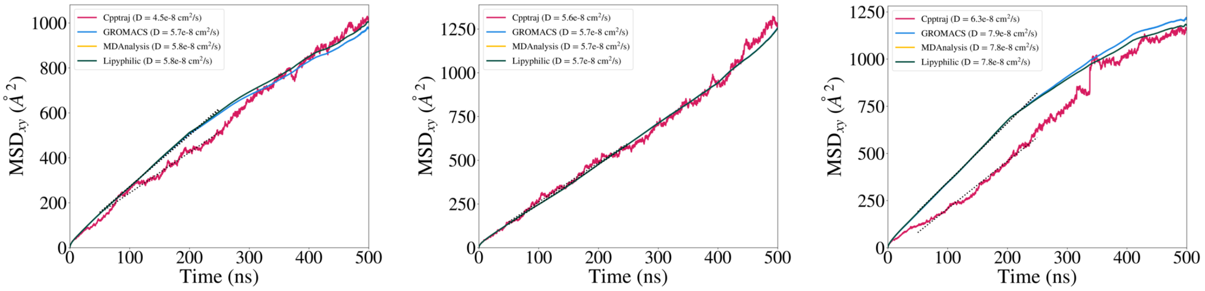


**Figure SI.4.** MSD comparison for the 256L-$j$W systems. Columns correspond to the hydration levels $j = 40, 80, 160$, from left to right.

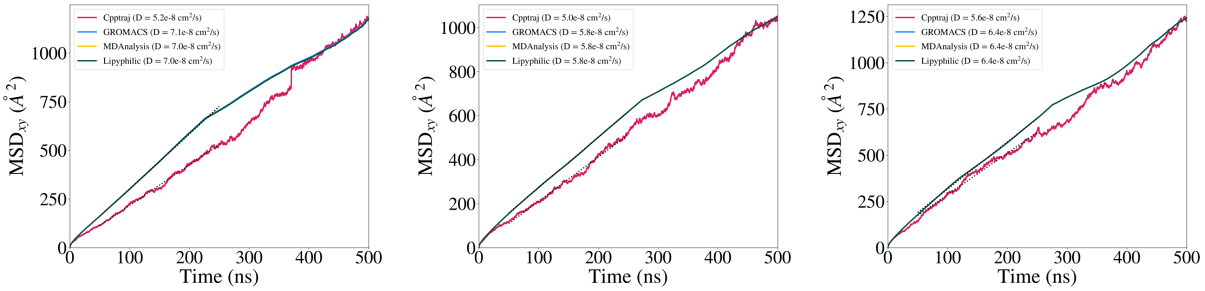


**Figure SI.5.** MSD comparison for the 512L-$j$W systems. Columns correspond to the hydration levels $j = 40, 80, 160$, from left to right.

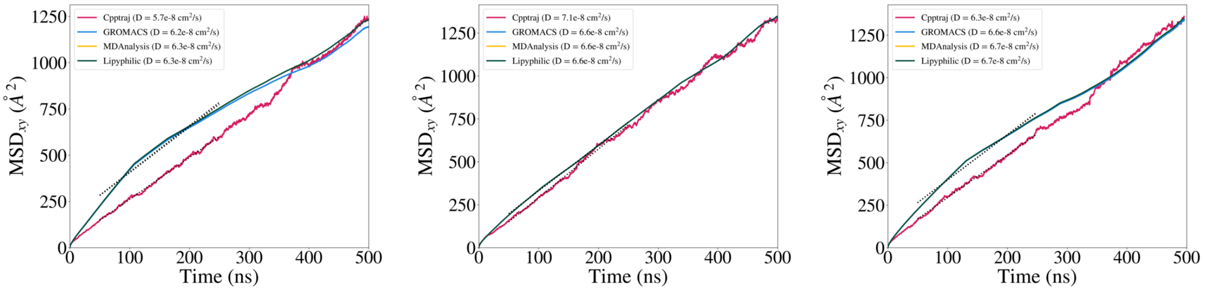


**Figure SI.6.** MSD comparison for the 1024L-$j$W systems. Columns correspond to the hydration levels $j = 40, 80, 160$, from left to right.

### Software comparison

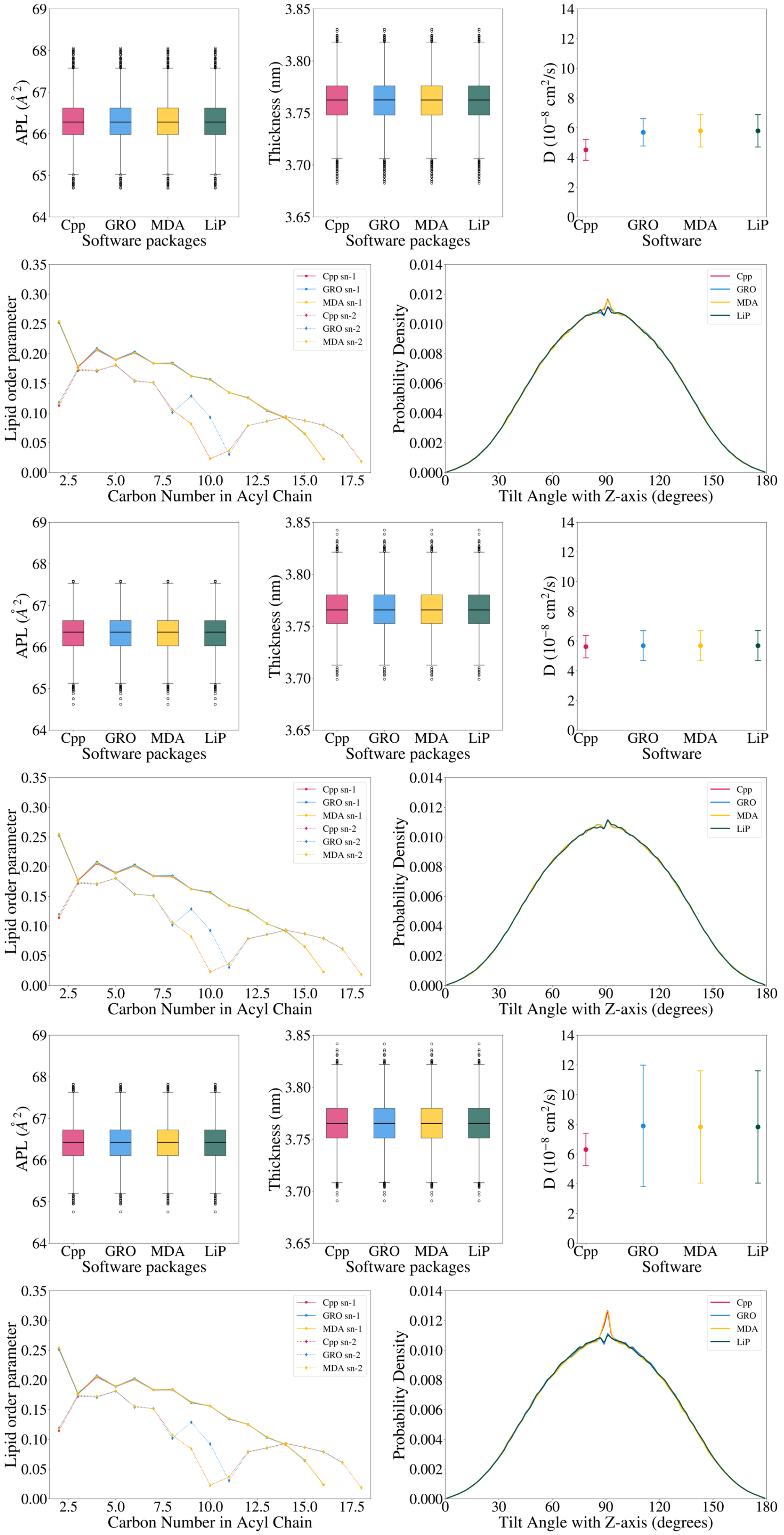


**Figure SI.7.** Software comparison for the 256L-$j$W systems. Each pair of rows corresponds to hydration levels $j = 40, 80, 160$, from top to bottom.

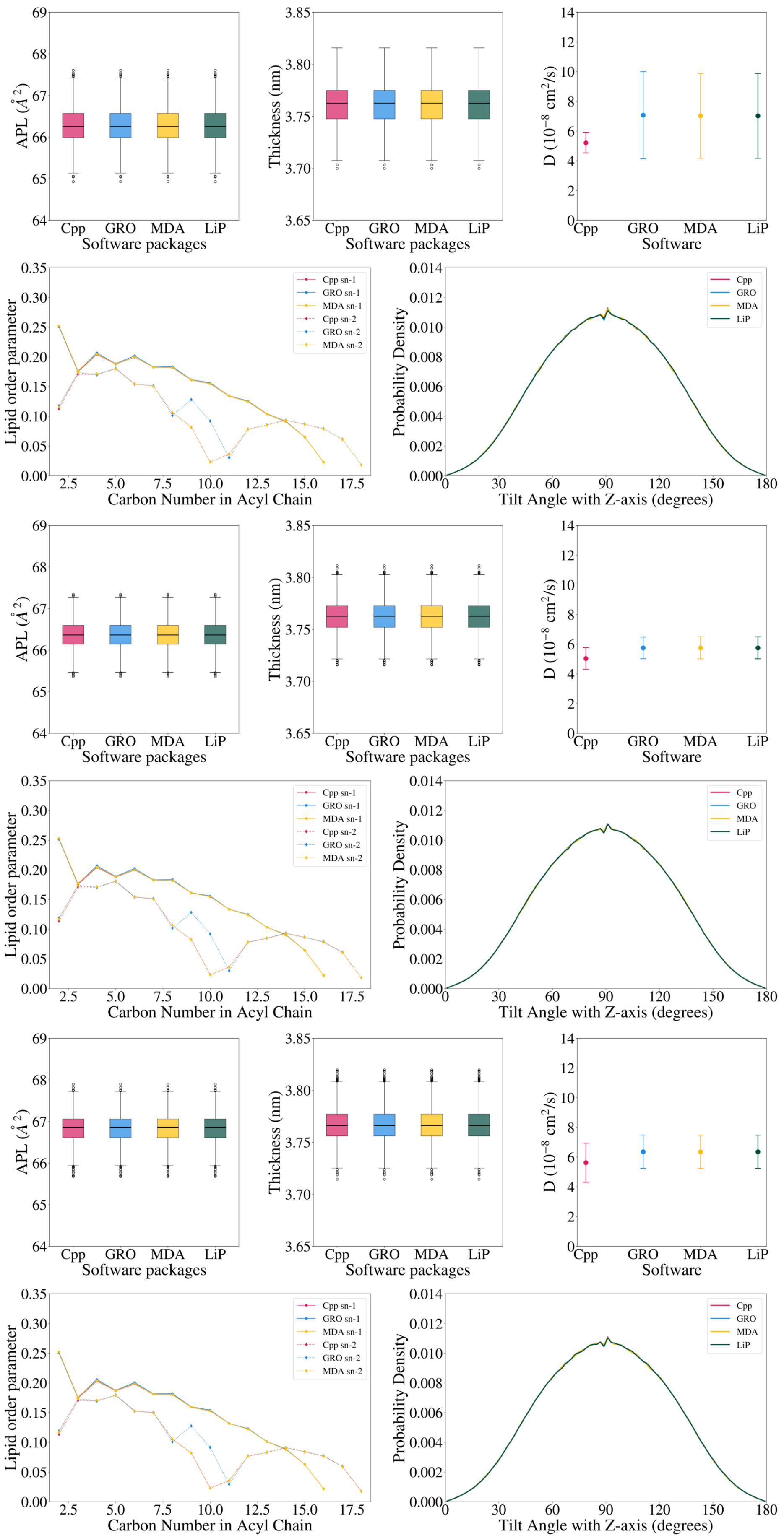


**Figure SI.8.** Software comparison for the 512L-$j$W systems. Each pair of rows corresponds to hydration levels $j = 40, 80, 160$, from top to bottom.

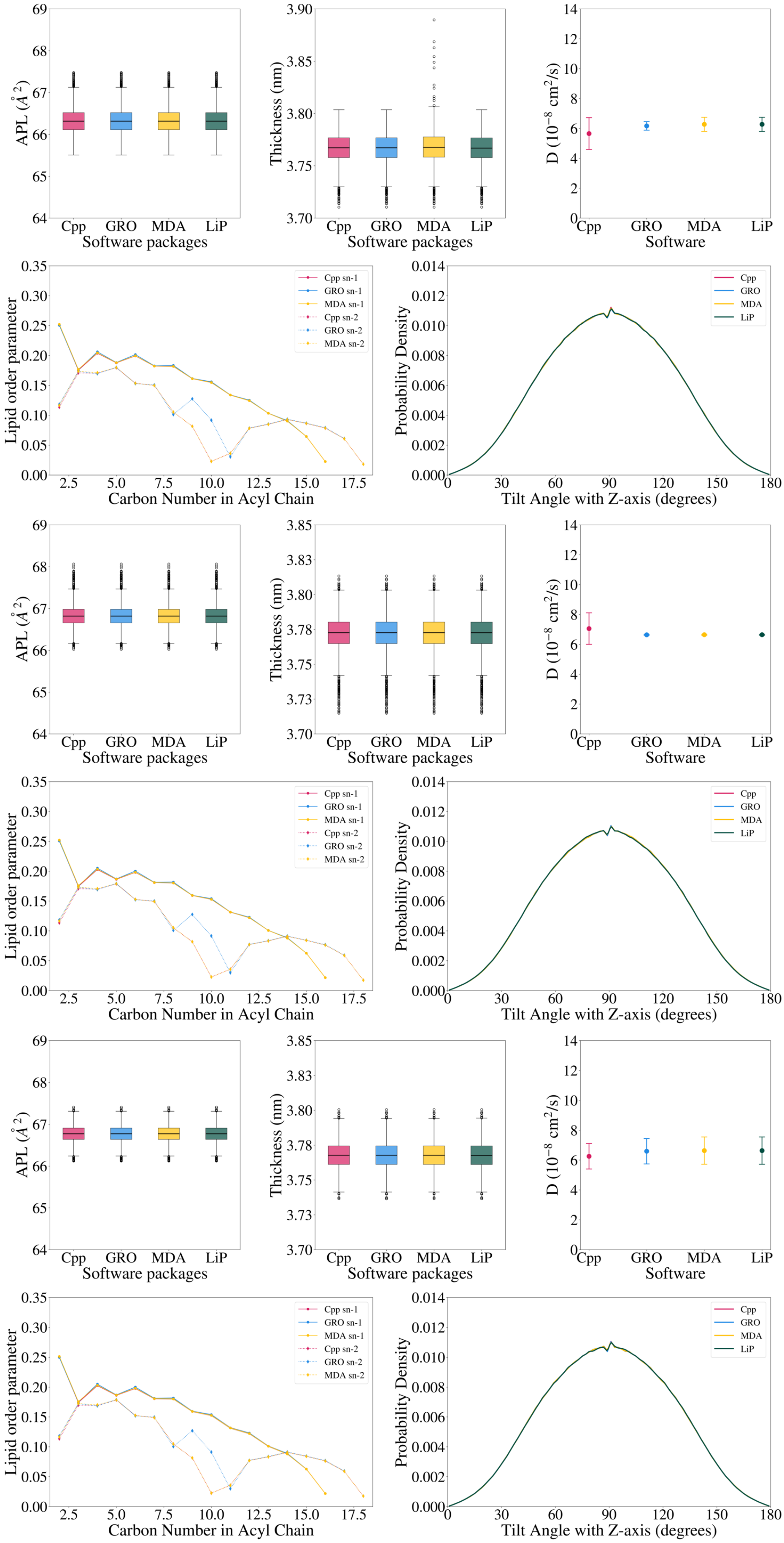


**Figure SI.9.** Software comparison for the 1024L-$j$W systems. Each pair of rows corresponds to hydration levels $j = 40, 80, 160$, from top to bottom.

### APL Voronoi Tesselation Analysis

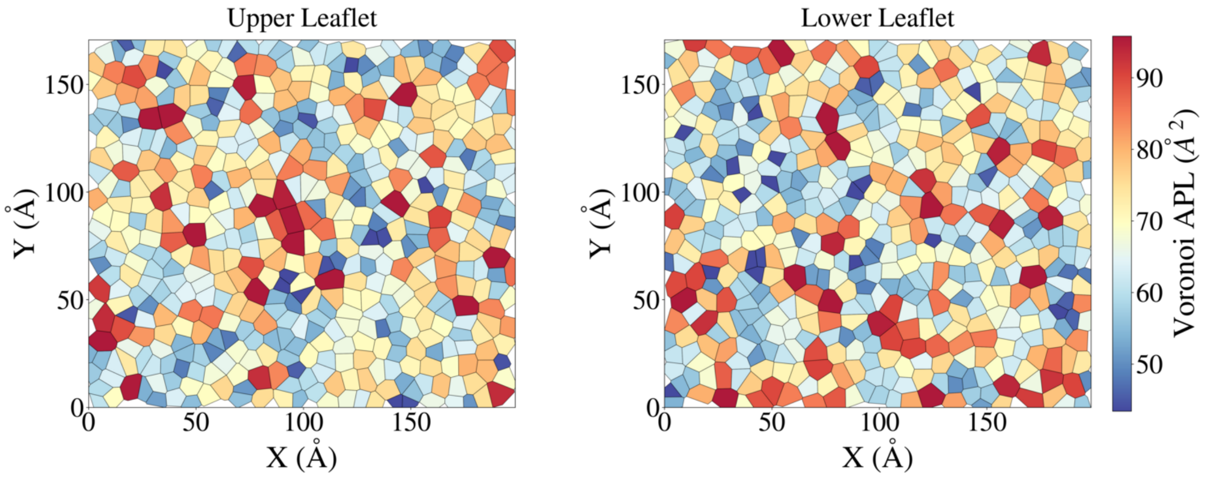


**Figure SI.10. 1024L-80W** Voronoi Tesselation map of both upper and lower leaflet.
